# High accuracy label-free classification of kinetic cell states from holographic cytometry

**DOI:** 10.1101/127449

**Authors:** Miroslav Hejna, Aparna Jorapur, Jun S. Song, Robert L. Judson

**Affiliations:** Department of Physics, University of Illinois, Urbana-Champaign; Carl R. Woese Institute for Genomic Biology, University of Illinois, Urbana-Champaign; Helen Diller Family Comprehensive Cancer Center, University of California, San Francisco; Department of Dermatology, University of California, San Francisco; Department of Bioengineering, University of Illinois, Urbana-Champaign

## Abstract

Digital holographic microscopy permits live and label-free visualization of adherent cells. Here we report the application of this approach for high accuracy kinetic quantitative cytometry. We identify twenty-six label-free optical and morphological features that are biologically independent. When used as a basis for machine learning, these features allow blind single cell classification with up to 95% accuracy. We present methods to control for inherent holographic noise, thereby establishing a set of reliable quantitative features. Together, these contributions permit continuous digital holographic cytometry for three or more days. Applying our approach to human melanoma cells treated with a panel of cancer therapeutics, we can track the response of each cell, simultaneously classifying multiple behaviors such as cell cycle length, motility, apoptosis, senescence, and heterogeneity of response to each therapeutic. Importantly, we demonstrate relationships between these phenotypes over time. This work thus provides an experimental and computational roadmap for low cost live-cell imaging and kinetic classification of heterogeneous adherent cell populations.

## Introduction

In cytometry, cells are classified into subpopulations of interest using measurable morphologic and molecular characteristics. Flow cytometry, the most universal method, is a powerful non-endpoint technique, but requires detachment of adherent cells, thus leading to loss of features associated with adhesion. The complementary approach of quantitative imaging with high content analysis, which allows for evaluation of adherent cells, often depends on reliable fluorescent labels for accurate classification of cell state ^1,2^. The cytotoxicity associated with fluorescent dyes and the phototoxicity associated with fluorescent light can limit the length of time single cells are tracked unperturbed ^3^. Additionally, reliable markers must be *a priori* identified to classify cell states of interest, despite observations that expression of single genes is often insufficient to predict cell state or behavior ^4^. With increasing demand for kinetic quantitative classification of subpopulations within heterogeneous cultures, there is a need for reliable label-free quantitative time-lapse solid-phase cytometry.

Phenotypic profiling is a high-content image analysis strategy for identifying cell state without the need for previously identified markers. The strategy uses computational reduction of a large number of general quantitative features to classify cells into distinct phenotypic profiles ^5,6^. This approach usually requires the fluorescent labeling of macromolecular markers, often DNA or histones. Label-free phenotypic profiling would permit both extended time-lapsed imaging (e.g. a two-week stem cell differentiation protocol) and cell state classification when fluorescent labeling is impractical or impossible (e.g. primary cell culture). There exist several methods for visualizing mammalian cells without the use of dyes or fluorescence ^7^. Some of these generate quantitative, instead of qualitative, data, allowing for the calculation of optical and morphological features that can be used to classify cells ^7–15^.

Recently, digital holographic microscopy (DHM) has emerged as one such quantitative method for visualizing adherent mammalian cell cultures ^16^. In DHM, one branch of a split laser beam passes through the transparent sample and recombines with the reference beam at an off-axis geometry, thereby generating interference ^15^. This interference pattern (the hologram) is used to reconstruct a wavefield of the illuminated cells, which can be visualized as a three-dimensional image ^17^. No light is absorbed by the cells, so DHM is non-phototoxic permitting long-term time-lapse imaging ^18^. Due to the relative affordability of commercially available DHM systems, this approach is becoming increasingly used for several applications, including cell counting, cell migration assays, monitoring for therapeutic resistance and motility characterization ^19^–^24^. In addition to morphological features derived from segmentation of DHM images, the interference pattern itself can be used to calculate absolute phase shift, providing an opportunity to determine cellular features related to optical density, such as thickness, texture, volume, and localized densities. Cell classification using these features is referred to as digital holographic cytometry (DHC). For example, DHC-derived volume can distinguish populations of pure X- or Y-chromosome-containing spermatozoa or populations enriched for cells in specific cell cycle states ^25,26^. However, with few exceptions, notably the identification of cells in M-phase of the cell cycle, the individual cells in these populations are rarely classified with single cell accuracy ^27–29^. In addition, the physical or biological meaning of optical features is dependent on technical, computational, and biological variables and must be interpreted with great care. For example, optical volume is known to be correlated with actual cell volume, cell detachment, cell flattening, calcium fluctuations, cell cycle, cell death, cell differentiation, and protein content ^17,30–37^. Finally, there is no established method for controlling the quality of a hologram in its application to DHC, making inter- and even intra-experimental comparison difficult. In order to be established as a reliable platform for quantitative cytometry, DHC must classify single cells with dependable high accuracy and provide consistent quantitative data that are comparable from one hologram to the next.

Here, we report our systematic characterization of hologram quality and the biological dependence structure of DHC-derived features. We developed methods both for aggregating features of unclear biological interpretation and for ensuring that the quantitative data derived from holograms are comparable within long-term kinetic experiments. We found that phenotypic profiling using twenty-six biologically independent features successfully classified multiple states with single cell resolution and a high degree of accuracy. We performed a time-lapsed screen of human melanoma cells treated with kinase inhibitors of known function to demonstrate the depth of information provided by this approach.

## Results

### Phenotypic profiling using DHC-derived features

Mammalian cell classification is complicated by a high level of cell heterogeneity, even in clonal cultures. We minimized biological variability by tracking individual cells over a time course and then determined their fate using established molecular markers. Once cells were grouped by verified state, we generated DHC-derived features from an earlier time point (24 hours), and investigated whether morphological features could accurately distinguish each state group (Fig. 1a). Holograms were captured using a HoloMonitor M4 DHC system ^38^ and acquired every hour to observe all cell divisions in the non-treated conditions. The holograms were processed computationally using HStudio to produce an intensity image representing a quantitative map of light-wave phase shifts ^38^–^40^. Individual cellular events were segmented and tracked from the resulting phase shift images. Homogeneous growth arrest and apoptosis were induced in the human melanoma cell line A375 with the CDK4/6 inhibitor PD0332991 and the kinase inhibitor Staurosporine, respectively (Fig. 1b) ^41^. PD0332991-treated cells failed to divide for ten days after treatment (Fig. 1c) and uniformly exhibited two markers of cellular senescence: B-galactosidase activity (Fig. 1d) and increased flow cytometric side scatter (Fig. 1e). Staurosporine-treated cells presented increased levels of cleaved CASP3 and PARP2 (Fig. 1f) and uniform surface expression of Annexin V (Fig. 1g), each being an established marker of pre-apoptosis. We analyzed the 24-hour images, and verified that each cell included in our analyses was either growth arrested (did not divide within 24 hours of time-point), pre-apoptotic (underwent cell death between 10 and 24 hours after time-point), or in M phase (divided into two cells within 1 hour after time-point) (Fig. 1h). We next asked whether commonly used DHC-derived features – perimeter, area, volume or thickness - could accurately classify each of these cell states (Fig. 1i-j and Fig. S1). Cellular features were calculated for each individual cell at the 24-hour time-point based upon the segmentation data and average refractive index. As previously reported, some feature distributions partially distinguished cell states. For example, area classified growth-arrested cells with 80% accuracy, and thickness separated pre-apoptotic cells from growth-arrested cells (Fig. 1i and Methods). However, no individual feature successfully separated all of the cell states (Fig. S1a). The combination of two or three features improved resolution to an average accuracy that plateaued around 68% (Fig. 1j-k, and Fig. S1b-c). Thus, the quantitative features derived from DHC can be sufficient for label free classification of some cell populations. However, single cell accuracy was not achieved for each of the verified homogeneous populations.

**Fig. 1:**
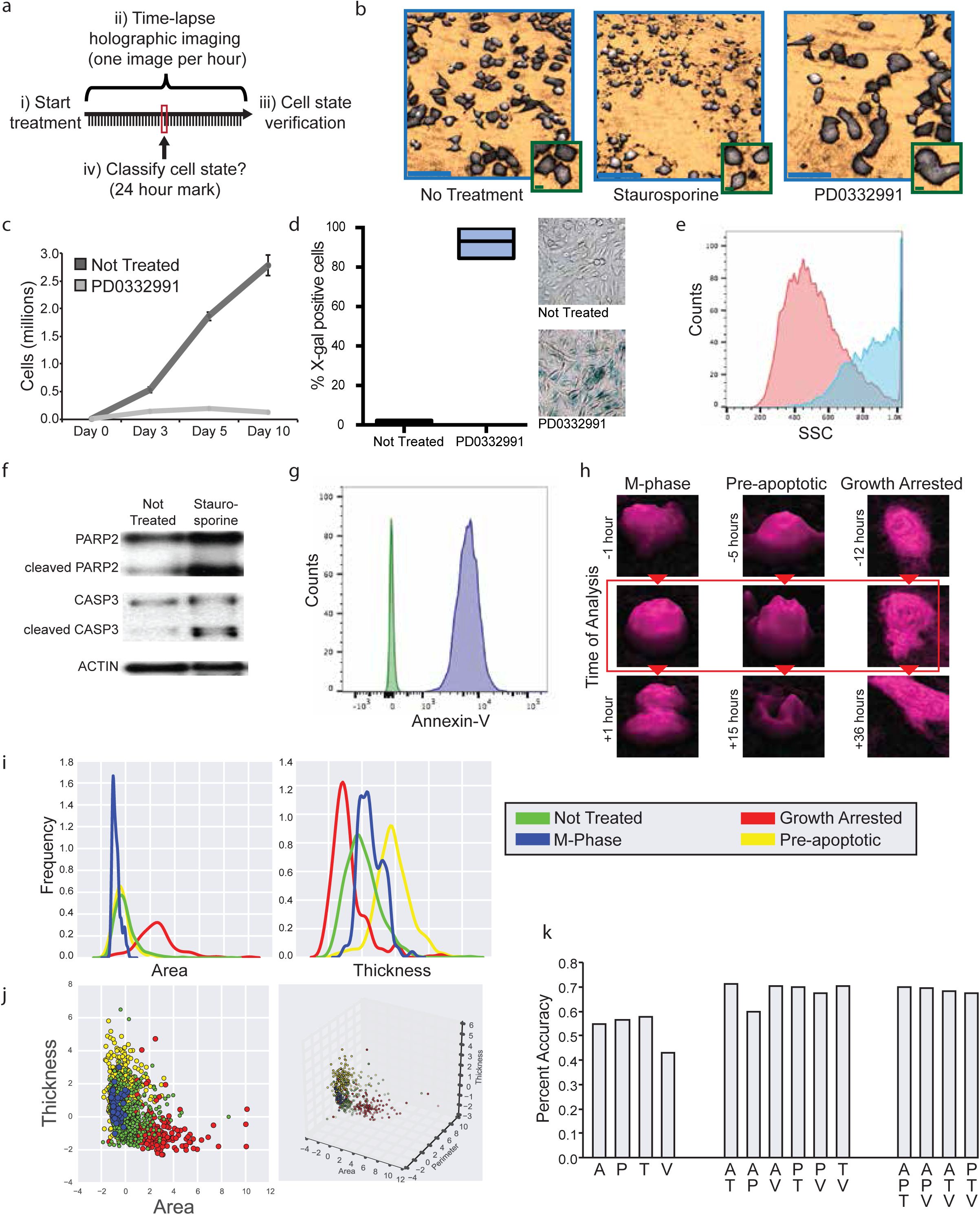
Classification of homogeneous populations with individual features. a) Strategy for DHC-based classification using cells of verified state b) Representative DHM images of indicated treatments with zoomed insets (scale bars: 100microns (blue), 10microns (green). c) Total number of cells for duration of experiment plus one week (n=4, standard deviation of mean). d) Representative images and quantification of X-gal positive cells after 7 days treatment (n=5, mean and range). e) Senescence-associated side scatter (SSC) after 7 days PD0332991 (blue) or control (red) treatment (representative of 4 experiments). f) Western blot of indicated proteins 7 days after indicated treatment (representative of 4 experiments). g) Annexin-V expression by flow analysis 1 day after Staurosporine (blue) or control (green) treatment (representative of 5 experiments). h) Representative images of cells designated as M-phase, pre-apoptotic, or growth arrested. Cell fates verified after the analyzed 24-hour time-point. i) Distribution of area and thickness for each cell state. j) 2D and 3D scatter plots of feature distribution for each cell state. k) Percent accuracy of cell classification using each set of one, two or three features: area (A), perimeter (P), thickness (T) or volume (V). Plots i-k used 470 pre-apoptotic, 195 growth arrested, 66 M-phase, and 1527 non-treated cells.

We reasoned that through increasing the number of independent features and conducting machine learning-based phenotypic profiling, we could better separate distinct cell states. As all features describing objects from holographic data are mathematically calculated from the phase shift profile, it is difficult to predict *a priori* which are biologically independent. To address this challenge empirically, we consolidated feature correlation data from thirty-five experiments covering a diverse cohort of morphologically distinct mouse and human cell populations - primary human melanocytes, human melanoma lines, and mouse mammary epithelial cells – each undergoing induced cell state transitions - including epithelial to mesenchymal transition (EMT), senescence, apoptosis, differentiation, and DNA-damage response (Fig. 2a). Each of the 42 optical, morphological, and positional features defined by HStudio were derived for each cell (Methods). As expected, many of the features were highly correlated, providing redundant information (Fig. 2b & Fig. S2). However, correlations were not always preserved across transitions (Fig. 2b). For example, cell flatness is highly correlated with cell length during EMT, but not during growth arrest. We therefore identified the minimal biological correlation for each pair of features, and combined those sets with conserved complete correlation to single metrics (Fig. 2c, Methods). The resulting set of 26 features provides biologically independent information that could be used to segregate distinct cell states. We applied a linear discriminant analysis (LDA) of the data set generated from the 24-hour holograms to project these 26 features onto the three-dimensional space that best separates the untreated, actively dividing, growth-arrested and apoptotic cell states. The resulting LDA plot clearly shows 4 clusters that can be separated by an unsupervised Gaussian Mixture Model without prior knowledge of the cell states (Fig. 3a-d). The four observed cell populations are classified with greater than 90% average accuracy, in some cases reaching 100% population separation. We then repeated the experiment ten additional times to generate independent test sets, without manual verification of individual cell state. These data sets contain greater biological heterogeneity, and represent a more realistic and tractable experimental set up for most applications. With these data sets, the clear separation into distinct clusters was not possible using only sets of one to three features, but the LDA predictions reliably increased the accuracy of cell classification by 22% to an average of 85% (Fig. 3e). These analyses demonstrate that DHC is a sufficient platform for reliable label-free single cell classification using machine-learning-based phenotypic profiling.

**Fig. 2:**
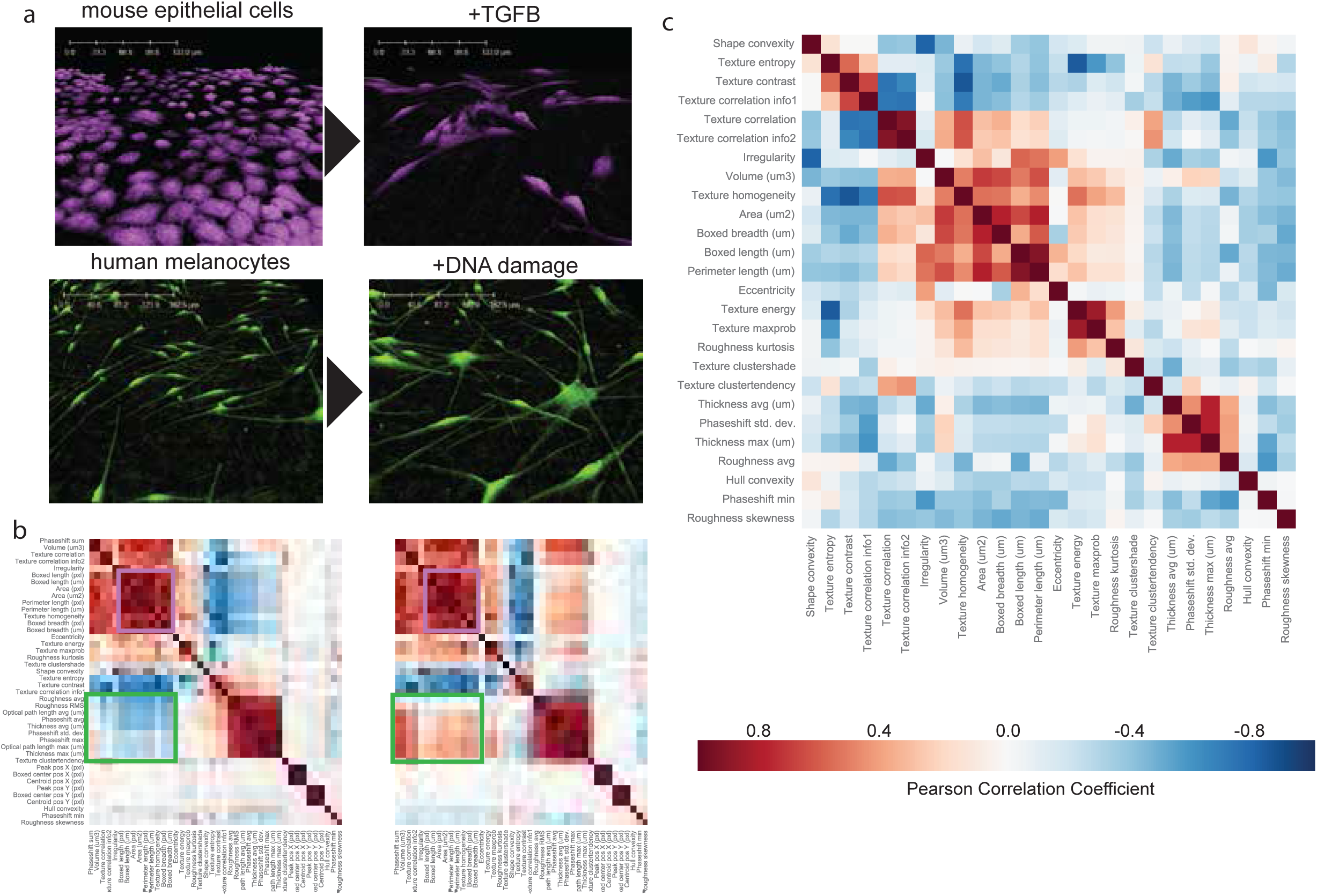
Identification of biologically independent features. a) Representative DHM images for two cell state transitions: EMT (top, NuMuG cells treated with TGFB) and DNA damage (primary human melanocytes treated with Doxorubicin) b) Two representative feature correlation matrices for mammary epithelial cells undergoing EMT (left) and human melanoma cells undergoing growth arrest (right). Areas of conserved (purple) or non-conserved (green) correlations are highlighted. c) Minimal correlation matrix from thirty-five experiments, each containing ˜1,000-10,000 cells (Fig. S2).

**Fig. 3:**
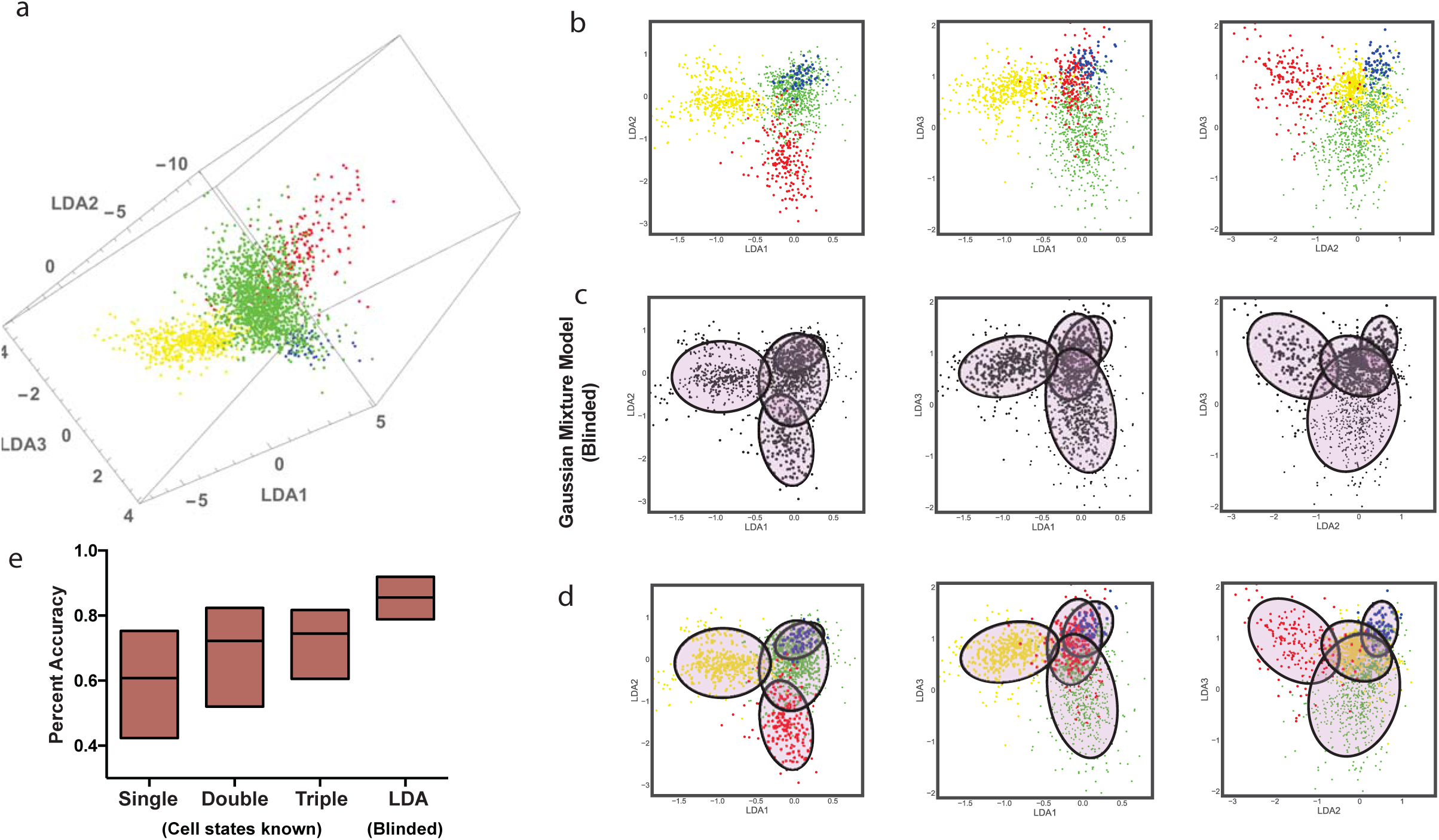
Phenotypic profiling using DHC-derived features. a-b) Three-dimensional LDA space derived using twenty-six features, demonstrating clusters of pre-apoptotic (yellow), growth-arrested (red), non-treated (green) or M-phase (blue) cells. Data use 470 pre-apoptotic, 195 growth arrested, 66 M-phase, and 1527 non-treated cells c) Four populations of cells identified by blinded Gaussian mix model of LDA space. Ellipses represent different views of three 3D predicted Gaussian distribution. d) Overlap of known cell states (dot colors) with predicted Gaussian distributions (ellipses). e) Distribution of percent accuracy of cell classification across all experiments using single, double or triple feature sets versus machine-learning based phenotypic profiling.

### Development of hologram quality control metrics for DHC

We next sought to characterize the accuracy of DHC classification for long-term time-lapse experiments using the aforementioned LDA model trained on the 24 hour data. We reasoned that imaging every hour and tracking individual cells provided two distinct advantages over other forms of cytometry. First, by running an LDA classification on the same cells every hour, errors in any individual classification (here, expected to be approximately 15% on average) could be reduced by serial re-classifications. Second, this approach would allow for the kinetic classification of individual cells – monitoring changes in cell state and its relationship to division and motility over time. We first verified that long-term imaging did not compromise the consistency of our holograms. As in any laser-based analytical method, the diodes widely used in DHM platforms require calibration prior to each experiment. When system calibration is suboptimal, visually comparable phase shift images can still be generated, but the underlying quantitative features differ in their intensity. This discrepancy can result in datasets with similar area-based features, but divergent thickness-based features from identical cells. From a classification perspective, this is similar to identical fields of fluorescent cells imaged with two different exposure times. Whereas such dissimilarities are easily distinguished in fluorescent-based imaging using background pixel intensity, no such metric exists for determining the compatibility of DHM images.

To determine the degree of thickness-based feature fluctuation over time in DHC, A375 cells were imaged every 24 hours for four days. To control for fluctuations in features related to biological differences, such as increased confluence, new passage- and density-matched cells were plated for each time-point. When data from at least eight holograms was consolidated, population means of thickness approached consensus across days and technical replicates (Fig. 4a-b, blue bars). However, when we compared thickness distributions across individual holograms within technical replicates – each generated from the same well within seconds - we observed notable population shifts (Fig 4c, blue and purple bars). For comparison, we repeated the experiments using a calibration that was suboptimal but still generated images that were visually comparable (Fig 4d). As expected, thickness-based features were more variable in these datasets (Fig 4b-c, red and orange bars). We sought to use this distribution of optimal and suboptimal datasets to define metrics of hologram quality control that would allow for the refining of feature data when conducting DHC-based single cell classification. We investigated whether traditional image quality control metrics such as signal-to-noise ratio (SNR) or average background signal (BgAv) (Fig. 4e) could bin holograms into groups of comparable accuracy, but found that BgAv did not correlate with the accuracy of classification (Fig. 4f) and SNR was not independent of cell state (Fig. 4g). Interestingly, we found that the degree of fluctuation of BgAv (BgSD) varied considerably between holograms (Fig. 4d-f,h). BgSD fluctuation was not influenced by cell state and inversely correlated with the classification accuracy (Fig. 4f-g). To determine whether categorization of holograms based upon BgSD increased the accuracy of classification, we removed holograms with BgSD > 0.5 (Fig. 4h, red line and Methods). Despite the consequential decrease in the number of data points, this filtering strategy selectively increased the classification accuracy in experiments containing suboptimal holograms by an average of 5%, but had no effect on a stability-calibrated system where most images passed the threshold (Fig. 4h-i). To determine how long a calibrated system continues to capture quantitatively comparable holograms during a time-lapse experiment, we collected 49 time-series, acquiring holograms and BgSD every hour for four days per experiment. As expected, BgSD drifted upward over time, but remained below the quality threshold for 40 hours (Fig. 4j). These data indicate that laser calibration is sufficient for at least four days of imaging and high accuracy LDA-based classification. We further identified BgSD as a reliable quality control metric for DHM images, which should be used to empirically determine the stability of a DHC system and the comparability of thickness-based features within and across datasets.

**Fig. 4:**
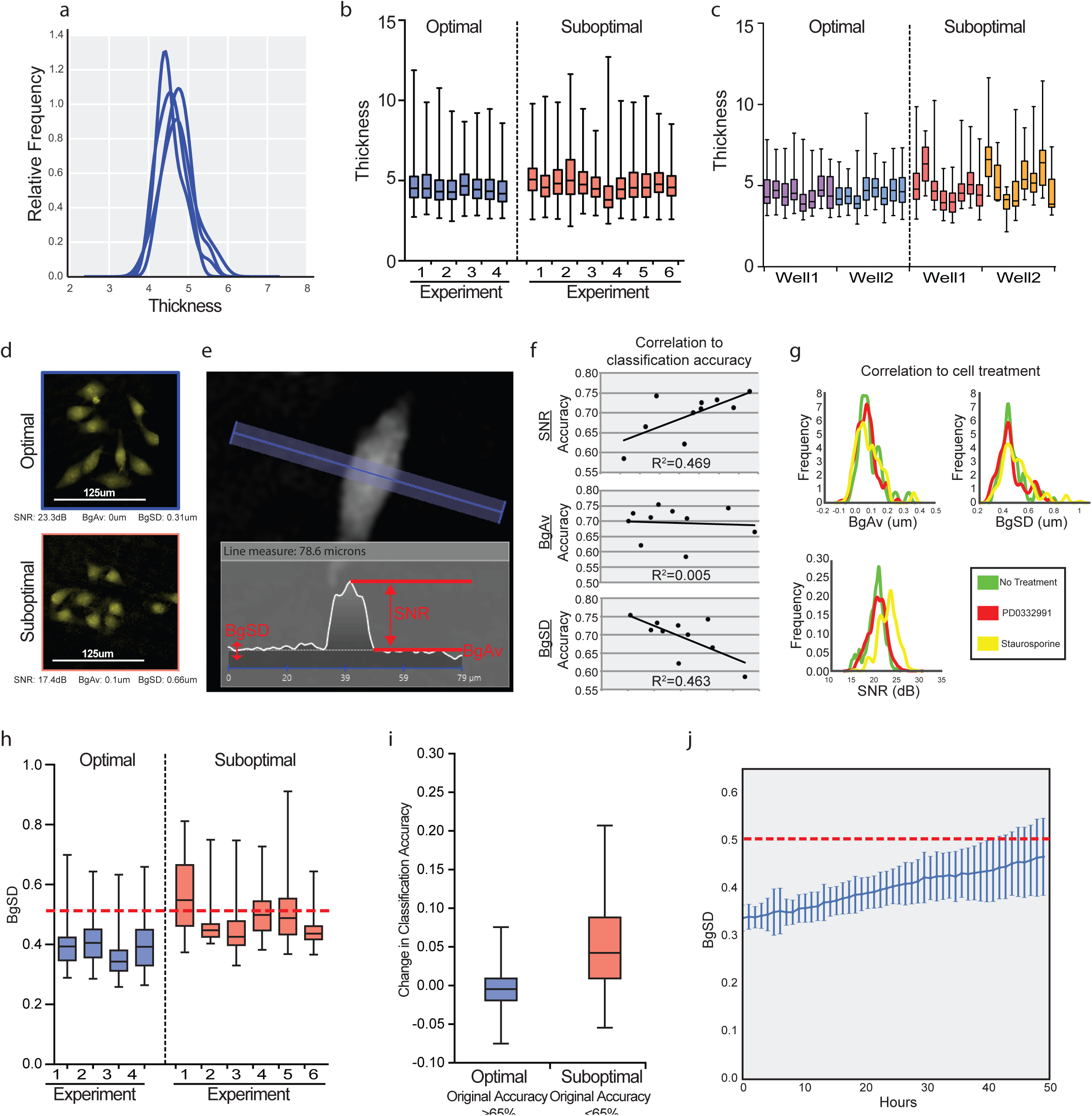
Development of hologram quality control metrics. a) Distribution of cellular thickness across four days and experiments. b) Distribution of cellular thickness from technical duplicates across ten experiments using optimally (blue) or suboptimally (red) calibrated diode. Each bar represents combined data from ten holograms c) Distribution of cellular thickness across individual holograms taken from the same well. d) Representative DHM images of cells taken with optimal or suboptimal calibration. e) Schematic of three potential hologram quality control (QC) metrics. Inset indicates intensity profile along blue line. f) Correlation of potential QC metrics (horizontal) with classification accuracy (vertical). g) Distribution of potential QC metrics segregated by cell state. h) BgSD for experiments from b-c. Dotted red line indicates QC threshold. i) Classification accuracy improvement after controlling for BgSD in experiments from b-c. j) BgSD over time from 49 image series taken continuously without calibration across 17 different experiments (standard deviation of mean). Dotted red line indicates QC threshold. Plots a-c and f-i are based on ˜24,000 cells across 10 experiments.

### Deep screening of kinase inhibitors

We next tested the capacity of our platform for deep screening by treating A375 human melanoma cells with a panel of well-characterized kinase inhibitors known to have toxic, growth-arresting or negligible effects. Growth curves were generated by counting the total number of cells in each series of phase shift images every four hours (Fig. 5a). On a population level, the effect of the compounds on cell proliferation matched previous observations. The targeted BRAF^V600E^ inhibitors Dabrafenib and Vemurafenib were acutely toxic along with the apoptosis-inducing compounds, Staurosporine and Doxorubicin. Other inhibitors of MAPK signaling were less toxic, whereas regulators of cell cycle slowed growth, and PI3K and BCR-ABL kinase inhibitors exhibited no population level effect. To investigate additional information provided by DHC, we monitored total cell movement, cell divisions, cell death and the 26 biologically independent features each hour for at least 40 cells per condition. Using the classifier we trained on our previous ten experiments (Fig. 3), we assigned a probability at each time point that a cell was pre-apoptotic or undergoing CDK4/6 inhibition. These probabilities, along with their relationships to cell division, cell death, and cell motility were visualized using “rocket” plots for each cell (Fig. 5b). We observed a clear association between cells predicted to be pre-apoptotic and eventual cell death. This state was associated with targeted BRAF^V600E^ inhibitors, apoptosis-inducing compounds, and MLN8347, consistent with the recent discoveries that AUORA kinase inhibitors are effective agents against BRAF^V600E^-driven melanoma cell lines (Fig. 5c) ^42,43^. The morphological state associated with CDK4/6 inhibition was cyclical in normal dividing cells, but sustained in growth-arrested cells. Interestingly, we observed substantial cell-to-cell heterogeneity in some, but not all treatments. We quantitated this heterogeneity by clustering the cell-level data contained within the rocket plots (Fig. 5c) and further clustered each treatment by apoptosis, arrest, mortality, and division rate (Fig. 5d and Methods). This provided a more detailed classification of the screened compounds than standard growth curves, successfully separating the targeted compounds from other acutely toxic compounds and compounds that induce growth arrest. Interestingly, the less acutely toxic compounds were grouped by compound vehicle used (either DMSO or PBS), indicating that DHC has the sensitivity to differentiate between the subtle effects of commonly used solvents within 48 hours of treatment. Within the DMSO cluster, inhibitors targeting the same kinase grouped together, demonstrating the platform can classify compounds by molecular function.

**Fig. 5:**
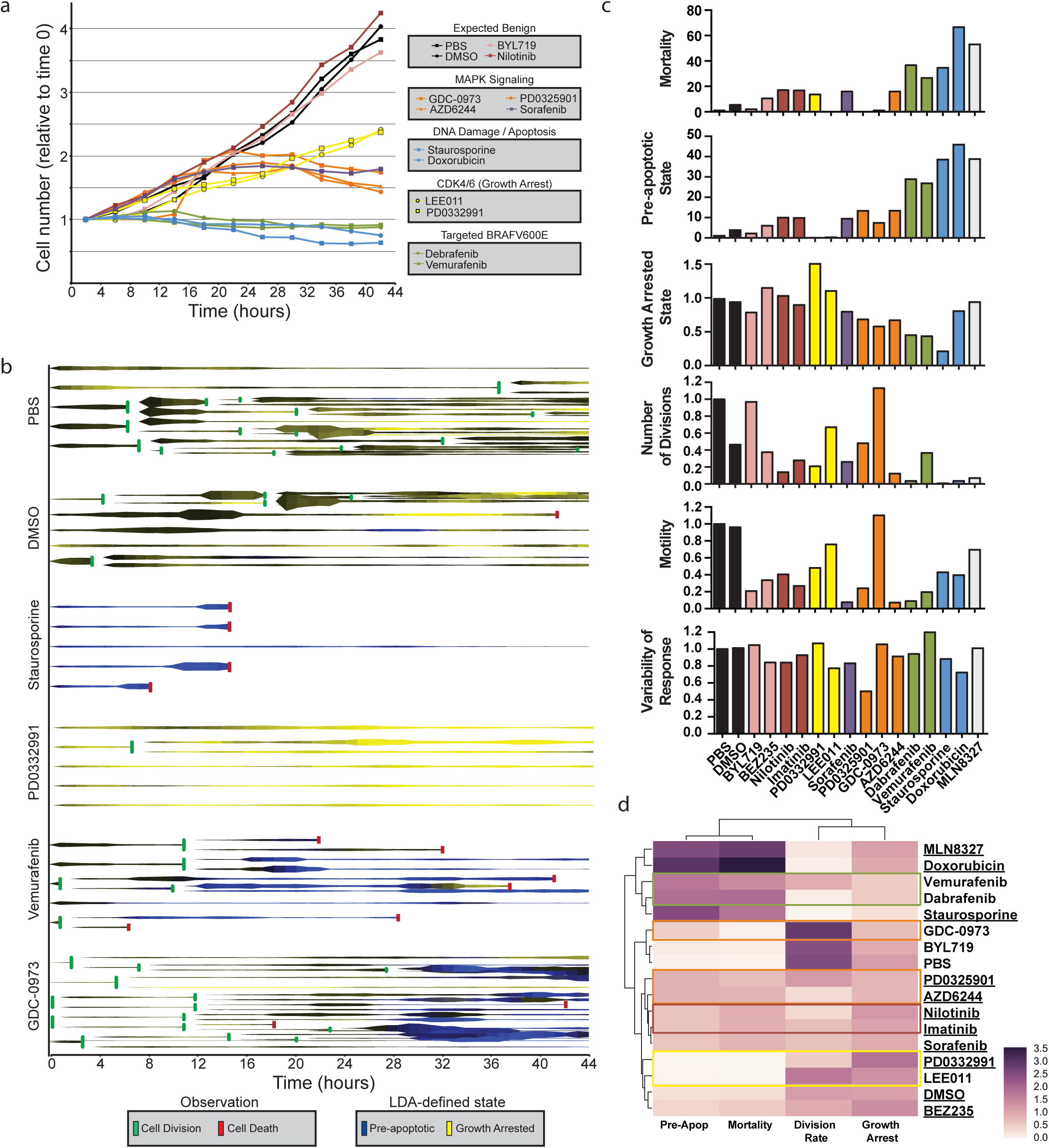
High depth screen of kinase inhibitors. a) Proliferation curves generated from time-lapse DHM-imaging, b) Example rocket plots of five representative cells over six conditions. Each horizontal line represents a single cell over time, where thickness correlates with microns moved since previous hour, colorimetric represents confidence of cell state based on morphology, and horizontal bars represent cell division or death. c) Cell apoptosis, arrest, division rate, motility, and variability of response relative to PBS for each compound. Colors represent compound classes as indicated in a). d) Heatmap depicting clustering of compounds by four super-metrics derived from rocket plots, normalized to the mean value of all conditions (see Methods). Targeted BRAF compounds (green), MEK inhibitors (orange), Bcr-Abl (red) and cell cycle inhibitors (yellow) are highlighted. Compounds dissolved in DMSO are underlined. Plots c-d are based on observation of 775 cells across 17 different treatment conditions.

## Discussion

### An accessible platform for label-free solid-phase time-lapse cytometry

Here we describe the development of a technical and computational pipeline for reliable and accessible DHC-derived label-free and kinetic classification of cell state. Previous pioneering reports have used similar platforms to compare DHM-derived features, but have been unable to achieve accurate single cell multi-state classification ^26^,^44^. By first establishing biologically homogeneous mammalian cell populations, we demonstrate that an extended set of features are amenable to machine-learning driven phenotypic profiling, increasing the accuracy of classification from a variable range of 11-89% to a more reliable range of 82-100%. Further, we developed methods for standardizing hologram quality for both experimental setup and longterm time-lapse imaging.

The basic technical set-up for DHM is relatively simple as compared to fluorescent microscopy and is consequently a more affordable option for phenotypic profiling. The system used in this study, the HoloMonitor M4, can be assembled inside a standard mammalian tissue culture incubator ^38^,^45^. DHM is inherently label-free and non-cytotoxic, allowing for long-term imaging of cell populations and tracking of individual cells within a population. As this analysis is non-endpoint, the cells can be used for a variety of purposes once the analysis is complete – for example, molecular verification of cell state as we did here, further propagation, or injection into a mouse.

We foresee a variety of uses for this platform. With the recent advent of CRISPR-Cas9 mediated genome engineering, the genomes of primary human cell lines can now be edited with high efficiency. This platform would allow for the direct comparison of edited cells to isogenic sibling control cells. Other uses include the identification and functional characterization of rare subpopulations of cells, or of transient cell populations within extensive differentiation protocols, without the need of known markers. It is important to note, as with all cytometry, that to accurately classify a cell into a biological state based upon its phenotypic profile, control populations of known state must be used to first characterize the optical morphological parameters of that biological state.

### High-depth screening

The strategy and platform described here provide particular advantages for small molecule screening. The major advantage is the depth of information gained with relatively few resources. In this study, all the data obtained over the entire time-period – proliferation, cell cycle length, motility, death, apoptosis, growth arrest and the relationship between each of these to each other and time – were acquired without operator intervention from a single treated well for each drug.

The presented data capture the heterogeneity of drug response between individual cells within each well. The method accurately identified high-potency general and targeted toxic compounds that reduced cell number through homogeneous induction of apoptosis, and distinguished these compounds from those that induced growth arrest or merely lengthened the cell cycle. Further, the analysis grouped compounds of similar molecular mechanism or solvent. Importantly, the morphological cell states analyzed in this screen - pre-apoptosis and CDK4/6-inhibition - were identifiable, because we were able to train the algorithm with homogeneous controls. In principle, any cell behavior associated with morphological changes could be similarly identified with proper controls. We expect our data analytics and visualization module to make this approach accessible to a wide range of studies.

## Methods

### Cell culture

The A375 human melanoma cell line was obtained from the Bastian lab (UCSF) and cultured in RPMI-1640 without HEPES supplemented with 10% FBS (Corning, 35-010-CV), 0.1mg/mL Pen-Strep, and 0.002mg/mL L-Glutamine. The cell line was validated based upon driver mutation (BRAF^V600E^ and CDKN2A) status. The NuMuMg mouse mammary cell line was obtained from the Derynk lab (UCSF) and cultured in DMEM H-21 supplemented with 10% FBS, 0.1mg/mL Pen-Strep, 0.002mg/mL L-Glutamine and 0.2mg/mL insulin (Sigma, 91077C). The cell line had the expected epithelial morphology and EMT response to TGFB, but is otherwise un-validated in our hands. Normal human melanocytes were purchased directly from ThermoFisher (C0245C), and cultured in Melanocyte medium (ThermoFisher, M254500) supplemented with HMGS (ThermoFisher, S0025). All lines in the laboratory, including those used during these studies, are verified routinely as mycoplasma negative using the ATCC Mycoplasma Detection Kit (30-1012-K).

For each holographic imaging experiment, passage-matched A375 cells were first split from wells of defined density (75% confluence) into 6-well culture plates (Sarsdedt, 83.3920) at 50,000 cells per well. NuMuMg cells and normal human melanocytes were seeded at 20,000 and 80,000 cells / well, respectively. After 24 hours, the media was replaced with 3mLs fresh media containing diluted small molecules and ligands at the below concentrations.

| Drug | Company | Cat. No. | Working conc |
| --- | --- | --- | --- |
| PD0332991 | Selleckchem | S1116 | 1000nM |
| Staurosporine | Tocris Bioscience | 1285 | 1nM |
| TGF-Beta | R&D Biosystem | 240-B-002 | 0.2ng/ml |
| Doxorubicin | Fischer | BP25161 | 10nM |
| Trametinib | Selleckchem | S2673 | 1nM |
| Cobimetinib | Selleckchem | S8041 | 0.84nM |
| Alpelisib | Selleckchem | S2814 | 100nM |
| Ribociclib | Selleckchem | S7440 | 61.4nM |
| Vemurafenib | Selleckchem | S1267 | 10nM |
| Dabrafenib | Selleckchem | S2807 | 0.16nM |
| Sorafenib | Selleckchem | S8599 | 10nM |
| PD0325901 | Selleckchem | S1036 | 0.1nM |
| MLN8327 | Selleckchem | S1133 | 100nM |
| Selumetinib | Selleckchem | S1008 | 2 nM |
| Imatinib | Selleckchem | S2475 | 200nM |
| Nilotinib | LC Labs | N-8207 | 10nM |
| PBS | UCSF-CCF |  | 0.01% of medium |
| DMSO | Sigma | 472301 | 0.01% of medium |

Aseptic optical lids (HoloLids, Phase Holographic Imaging) were placed on each plate to prevent condensation and surface vibrations to allow for 36 to 52 hours of time-lapse imaging. For cell state verification, cells were either: i) stained using a Senescence B-Galactosidase kit (Cell Signaling, 9860) and imaged using a Lieca Dmi1 light microscope; ii) analyzed for changes in side-scatter using a BD FACSCalibur; iii) lysed for Western analysis using RIPA Lysis and Extraction Buffer (ThermoFisher 89900) supplemented with Halt Protease Inhibitor cocktail (ThermoFisher 87786); or iv) stained with a Pacific Blue tagged Annexin V antibody (Biolegend 640917) for 15minutes before FACS analysis.

### Molecular Biology

Protein extracts were mixed with NuPAGE LDS Sample Buffer (ThermoFisher NP0007), NuPAGE Sample Reducing Agent (ThermoFisher NP0004) and heated for 10min at 70°C. Protein (10ug/lane) was resolved on NuPAGE Novex 4-12% Bis-Tris Protein Gels, 1.0 mm, 15-well (ThermoFisher NP0323BOX) and transferred using a Mini-Trans-Blot Cell onto 0.22Um PVDF membranes (ThermoFisher 88520). Membranes were blocked with Membrane Blocking Solution (ThermoFisher 000105) for 1 hour at room temperature (22°C). The membranes were then incubated overnight at 4°C with primary antibodies at the following dilutions: anti-Caspase 3 (CST 9661) 1:1000, anti-PARP2 (CST 9542) 1:1000, anti-Actin (CST 4970) 1:5000. The membranes were then washed four times with PBS and 0.5% Tween20 (TBST) and incubated with horseradish peroxidase-conjugated secondary antibody (anti-rabbit NA934) 1:15000 for 1 h at room temperature. The membranes were then washed four times with TBST and were visualized with Lumina Forte Western HRP substrate (Millipore WBLUF0500).

### Digital holographic microscopy and analysis

Digital holographic microscopy was performed using a HoloMonitor M4 imaging cytometer with high precision automated stage (Phase Holographic Imaging, Lund, Sweden) in which the object beam and reference beam of 50:50 split 635nm laser light are each, respectively, either passed through the sample then collected by a PLN 20x objective (Olympus) or tilted to create an off-axis configuration, before being directed to a 1.3MP CMOS global shutter USB 2.0 camera ^38^. Wavefields were reconstructed by HStudio using a combination of the basic Fesnel transform and angular spectrum methods, as previously reported ^39^,^40^. Absolute thickness was estimated by approximating and holding constant the average difference in the refractive index between cells and media as 0.04.

Holograms of A375 cells were generated in a standard mammalian cell culture incubator at 37°C and 5% CO_2_. DHM images were generated and segmented using the HStudio (v2.6.3). The software calculates the most well focused image from the range of potential focal distances. DHM images were uniformly segmented using an Otsu’s method algorithm with an optical thickness threshold of 130 and minimal object size of 26. Optical metrics in the non-segmented space (Fig. 4) were calculated as follows:

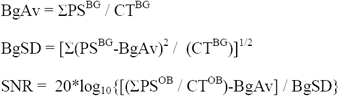

Where PS is the phase shift value of a pixel. CT is the pixel count. Superscript BG refers to all pixels not included in segmented objects. Superscript OB refers to all pixels included in segmented objects.

### Correlation matrices

Pearson correlation coefficient was used to assess the biological independence of each pair of features generated by HStudio. Since correlation between features depends on the biological state of cells and the induced transition (Fig. S2), the smallest correlation coefficient (in absolute size) among 35 experiments measuring cell transitions between different states under different treatments was plotted in Fig. 3c to evaluate the biological independence of each feature pair. Of 42 variable cell metrics provided by HStudio, we eliminated features that have more than 98% correlation in all 35 experiments. We also eliminated features that were a constant multiple of another feature already used and metrics that measure position of a cell within the image. The resulting list of non-redundant features used for classification of cell states is as follows: Area (μm^2^), Boxed breadth (μm), Boxed length (μm), Eccentricity, Hull convexity, Irregularity, Perimeter length (μm), Phaseshift min, Phaseshift std. dev., Roughness avg, Roughness kurtosis, Roughness skewness, Shape convexity, Texture clustershade, Texture clustertendency, Texture contrast, Texture correlation, Texture correlation info1, Texture correlation info2, Texture energy, Texture entropy, Texture homogeneity, Texture maxprob, Thickness avg (μm), Thickness max (μm), Volume (μm^3^).

### Cell state classification

Linear discriminant analysis was used to project the cell features into a two or three-dimensional linear subspace that separates the different treatment categories of cell states (Fig. 3a-b). A Gaussian mixture model was then fitted to the projected states in order to assign class probabilities to regions corresponding to the different cell states (Fig. 3c-d). We bootstrap sampled 1000 cells from each treatment group in order to have cell state clusters of comparable size in the training set.

When considering only commonly used DHC-derived features (T, P, A, or V), clusters corresponding to different cell states were poorly separated; thus, unbiased Gaussian mixture model fitting did not yield a useful classification in this case. Instead, in order to assess the most favorable classification accuracy under Gaussian models, we fitted a Gaussian model to each condition separately (supervised learning) and used the resulting mixture model to predict cell states (Fig. 1j, 3e, and 4f-g, i).

### Hologram Filtering

To determine whether removing the holograms with high values of BgSd improved the classification accuracy, incrementally increasing values of the thresholds (0.45,0.5,0.55,0.6) were set and the average classification accuracy across ten replicate experiments (4 with optimal calibration and 6 with suboptimal calibration) were evaluated (Fig. 4h-i). The optimal threshold value of BgSd that lead to the highest average classification accuracy is BgStd=0.5. With suboptimal calibration, filtering the holograms lead to the average removal of 40% of holograms and 5% average increase in classification accuracy. With optimal calibration, on the other hand, only 10% of holograms were removed and the increase in classification accuracy was less than 1%.

### Single cell analysis

Each cell was tracked throughout a 48h time frame with measurement performed every hour. At each time point, the cell state classification model was used to assign to each cell a probability of three states (growth arrested (CDKi), pre-apoptotic (St) or neither). For display purposes, probabilities were smoothed with Savitzky-Golay filter with degree 2 and window 5 (Fig. 5b). In order to better understand the overall response of cells to treatment, cell behavior was condensed into 6 biologically relevant super-metrics for population analysis: pre-apoptotic indicator, growth arrested indicator, division rate, mortality, motility, and variability of response (Fig. 5c). The pre-apoptotic indicator is the probability of a cell within the population being in the St state and set to 1 upon cell death. Growth arrested indicator is the probability of a cell within the population being in the CDKi state. Division rate is the probability of a cell within the population dividing. Mortality is the ratio of dead to all cells at the last time point. Motility is the probability that a cell within a population is undergoing a period of hypermotility. Variability of response is defined as the average variance of the predicted morphology-defined state, division rate and motility rate between the different cells in the experiment. We normalized the super-metrics to have the average value across all cell conditions to be 1. Hierarchical clustering of the drugs was performed using Euclidean distance and average linkage (Fig. 5d).

## Code availability

A python code used for our analysis is available upon request.

## Acknowledgments

This research was supported in part by NIH R01CA163336 and the Grainger Engineering Breakthroughs Initiative to J.S.S. and by NIH DP5OD019787 and the Sandler Foundation and Program for Breakthrough Biomedical Research to R.L.J. We thank E. Markegard and M. McMahon for providing kinase inhibitors, B. Bastian for the A375 cell line, and E. Holden and L. Weinberger for critical reading of the manuscript.

## Author Contributions

A.J. performed all cellular and molecular experiments and all holographic imaging and hologram processing. M.H. wrote all code and conducted all computational experiments including correlation matrices, cell classification, machine learning, and single cell analysis. J.S.S. and R.L.J. conceived of the experiments. M.H., A.J., J.S.S, and R.L.J. wrote the manuscript.

## References

1. Henriksen, M., Miller, B., Newmark, J., Al-Kofahi, Y. & Holden, E. Recent Advances in Cytometry, Part A - Instrumentation, Methods. Methods in Cell Biology 102, (2011).

2. Taylor, D., Haskins, J. & Giuliano, K. High content screening: A powerful approach to systems cell biology and drug discovery. Humana Press (2007).

3. Tinev, J.-Y. et al. A quantitative method for measuring phototoxicity of a live cell imaging microscope. Methods in enzymology 506, 291–309 (2012).

4. Trapnell, C. Defining cell types and states with single-cell genomics. Genome Research 25, 1491–1498 (2015).

5. Neumann, B. et al. Phenotypic profiling of the human genome by time-lapse microscopy reveals cell division genes. Nature 464, 721–727 (2010).

6. Bougen-Zhukov, N., Loh, S., Lee, H. & Loo, L. Large-scale image-based screening and profiling of cellular phenotypes. Cytometry Part A 91, 115–125 (2016).

7. Mann, C. J., Yu, L., Lo, C.-M. & Kim, M. K. High-resolution quantitative phase-contrast microscopy by digital holography. Optics Express 13, 8693 (2005).

8. Zangle, T. A. & Teitell, M. A. Live-cell mass profiling: an emerging approach in quantitative biophysics. Nat. Methods 11, 1221–8 (2014).

9. Kasprowicz, R., Suman, R. & O’Toole, P. Characterising live cell behaviour: Traditional label-free and quantitative phase imaging approaches. The International Journal of Biochemistry & Cell Biology 84, 89–95 (2017).

10. Mir, M., Bhaduri, B., Wang, R., Zhu, R. & Popescu, G. Quantitative phase imaging. Progress in Optics 57, 133–211 (2012).

11. Masters, B. Quantitative Phase Imaging of Cells and Tissues. Journal of Biomedical Optics 17, 029901–029902 (2012).

12. Tian, L. & Waller, L. Quantitative differential phase contrast imaging in an LED array microscope. Optics Express 23, 11394–11403 (2015).

13. Marquet, P., Rappaz, B. & Pavillon, N. New Techniques in Digital Holography: Chapter 5 Quantitative Phase-Digital Holographic Microscopy: A New Modality for Live Cell Imaging. 169–217 (Wiley, 2015). doi:10.1002/9781119091745.ch5

14. Majeed, H. et al. Quantitative phase imaging for medical diagnosis. Journal of Biophotonics 10, 177–205 (2017).

15. Marquet, P. et al. Digital holographic microscopy: a noninvasive contrast imaging technique allowing quantitative visualization of living cells with subwavelength axial accuracy. Optics Letters 30, 468–470 (2005).

16. Alm, K. et al. Digital Holography and Cell Studies, Holography, Research and Technologies: Chapter 11 Digital holography and cell studies. InTech 237–252 (2011). doi: 10.5772/15364

17. Shaked, N. T., Satterwhite, L. L., Rineheart, M. T. & Wax, A. Quantitative Analysis of Biological Cells Using Digital Holographic Microscopy. (InTech, 2011). doi:10.5772/15122

18. Rappaz, B., Kuttler, F., Breton, B. & Turcatti, G. Digital Holographic Imaging for Label-Free Phenotypic Profiling, Cytotoxicity, and Chloride Channels Target Screening. Methods in Pharmacology and Toxicology 307–325 (2015). doi:10.1007/978-1-4939-2617-6_17

19. Mölder, A. et al. Non-invasive, label-free cell counting and quantitative analysis of adherent cells using digital holography. Journal of microscopy 232, 240–7 (2008).

20. Sun, H. et al. Visualization of fast-moving cells in vivo using digital holographic video microscopy. Journal of Biomedical Optics 13, (2008).

21. Zhang, Y. et al. Reversal of Chemoresistance in Ovarian Cancer by Co-Delivery of a P-Glycoprotein Inhibitor and Paclitaxel in a Liposomal Platform. Molecular Cancer Therapeutics 15, 2282–2293 (2016).

22. Gao, Y. et al. Loss of ERα induces amoeboid-like migration of breast cancer cells by downregulating vinculin. Nature Communications 8, 14483 (2017).

23. Persson, H., Li, Z., Tegenfeldt, J. & Oredsson, S. From immobilized cells to motile cells on a bed-of-nails: effects of vertical nanowire array density on cell behaviour. Scientific … (2015).

24. Persson, J., Mölder, A., Pettersson, S. & Alm, K. Cell motility studies using digital holographic microscopy. Microscopy: Science, Technology, Applications and Education 1063–1072 (2010).

25. Miniotis, M., Mukwaya, A. & Wingren, A. G. Digital holographic microscopy for noninvasive monitoring of cell cycle arrest in L929 cells. PloS one 9, e106546 (2014).

26. Merola, F., Miccio, L., Memmolo, P., Caprio, D. G. & Galli, A. Digital holography as a method for 3D imaging and estimating the biovolume of motile cells. Lab Chip 4512–4516 (2013). doi:10.1039/c3lc50515d

27. Luther, E. & Kamentsky, L. Resolution of mitotic cells using laser scanning cytometry. Cytometry 23, 272–278 (1996).

28. Kemper, B., Bauwens, A., Vollmer, A., Muthing, J. & Bally, G. von. Label-free quantitative cell division monitoring of endothelial cells by digital holographic microscopy. Journal of Biomedical Optics 15, 036009–1–6 (2010).

29. Mir, M. et al. Optical measurement of cycle-dependent cell growth. Proceedings of the National Academy of Sciences 108, 13124–13129 (2011).

30. Kemmler, M. et al. Noninvasive time-dependent cytometry monitoring by digital holography. Journal of Biomedical Optics 12, 064002–1–10 (2007).

31. Schnekenburger, J. et al. Digital holographic imaging of dynamic cytoskeleton changes. Medical Laser Application 22, 165–172 (2007).

32. Pavillon, N. et al. Cell morphology and intracellular ionic homeostasis explored with a multimodal approach combining epifluorescence and digital holographic microscopy. Journal of Biophotonics 3, 432–436 (2010).

33. Pavillon, N. et al. Early cell death detection with digital holographic microscopy. PLoS One 7, (2012).

34. Alm, K. et al. Cells and Holograms–Holograms and Digital Holographic Microscopy as a Tool to Study the Morphology of Living Cells. InTech (2013). doi:10.5772/54505

35. Zlotek-Zlotkiewicz, E., Monnier, S., Cappello, G., Berre, M. & Piel, M. Optical volume and mass measurements show that mammalian cells swell during mitosis. The Journal of Cell Biology 211, 765–774 (2015).

36. Chalut, K. J., Ekpenyong, A. E., Clegg, W. L., Melhuish, I. C. & Guck, J. Quantifying cellular differentiation by physical phenotype using digital holographic microscopy. Integr Biol (Camb) 4, 280–4 (2012).

37. El-Schich, Z et al. Induction of morphological changes in death-induced cancer cells monitored by holographic microscopy. Journal of Structural Biology 189, 207–212 (2015).

38. Sebesta, M. et al. HoloMonitor M4: holographic imaging cytometer for real-time kinetic label-free live-cell analysis of adherent cells. Proc. SPIE 9718, Quantitative Phase Imaging II (2016). doi:10.1117/12.2216731

39. Kim, M., Yu, L. & Mann, C. Interference techniques in digital holography. Journal of Optics A: Pure and Applied Optics 8, S518 (2006).

40. Cuche, E., Marquet, P. & Depeursinge, C. Simultaneous amplitude-contrast and quantitative phase-contrast microscopy by numerical reconstruction of Fresnel off-axis holograms. Applied optics 38, 6994 (1999).

41. Fry, D. W. et al. Specific inhibition of cyclin-dependent kinase 4/6 by PD 0332991 and associated antitumor activity in human tumor xenografts. Molecular cancer therapeutics 3, 1427–38 (2004).

42. Xie, L. & Meyskens, F. The pan-Aurora kinase inhibitor, PHA-739358, induces apoptosis and inhibits migration in melanoma cell lines. Melanoma research 23, 102–13 (2013).

43. Porcelli, L. et al. Aurora kinase B inhibition reduces the proliferation of metastatic melanoma cells and enhances the response to chemotherapy. Journal of Translational Medicine 13, (2015).

44. Bettenworth, D. et al. Quantitative Stain-Free and Continuous Multimodal Monitoring of Wound Healing In Vitro with Digital Holographic Microscopy. PLoS ONE 9, e107317 (2014).

45. Peter, B. et al. Incubator proof miniaturized Holomonitor to in situ monitor cancer cells exposed to green tea polyphenol and preosteoblast cells adhering on nanostructured titanate surfaces: validity of the measured parameters and their corrections. Journal of biomedical optics 20, 067002 (2015).

